# MTHFR gene A1298C polymorphism and Alzheimer’s disease susceptibility

**DOI:** 10.1101/785063

**Authors:** Vandana Rai

## Abstract

Methylenetetrahydrofolate reductase (MTHFR) is a crucial enzyme involved in homocysteine/methionone metabolism. It catalyzes the conversion of 5,10methlenetetrahydrofolate in to 5methyltetrahydrofolate. A number of studies have examined the association of MTHFR A1298C polymorphism as risk factor for Alzheimer’s disease (AD), but the results were contradictory. To clarify the influence of MTHFR A1298C polymorphism on Alzheimer’s disease (AD), a meta-analysis of ten case-control studies was carried out. Four electronic databases were searched up to August, 2019 for suitable articles. The pooled odds ratios (ORs) with 95% confidence intervals (95% CIs) were used to evaluate the association. All statistical analyses were performed by MetaAnalyst program.

The results of meta-analysis suggested that except allele contrast model, A1298C polymorphism is not risk for Alzheimer’s disease using overall comparisons in three genetic models (C vs. A: OR= 1.26, 95%CI= 0.912-1.76, p= 0.04; CC+AC vs. AA: OR= 1.43; 95%CI= 0.85-2.44; p=0.05; CC vs. AA: OR= 1.16, 95%CI= .88-1.55, p= 0.51; AC vs. AA: 1.55; 95%CI= 0.81-2.93,p=0.07). Publication bias was absent in all five genetic models. In conclusion, results of present meta-analysis showed no significant association between MTHFR A1298C polymorphism and AD risk.

## Introduction

Alzheimer’s disease (AD) is one of the major neurodegenerative diseases in elderly population. It is the most common form of dementia, affecting 1 in 8 individuals older than 60 years of age. Most AD cases are late in onset and are probably influenced by both genetic and environmental factors. Epidemiological studies have demonstrated that elevated levels of plasma homocysteine (Hcy) may play an important role in the pathogenesis of AD [1-3].

Folic acid/folate is essential for cellular methylation, DNA synthesis, and homocysteine metabolism. MTHFR and methionine synthase reductase (MTRR) are two important enzymes of folate pathway and dysfunction of these genes increases plasma homocysteine concentration [4,55). Both genes show polymorphism as MTHFR C677T, A1298C [6-9] and MTRR A66G [10-12] and frequency of these polymorphisms varies greatly word wide. MTHFR enzyme required for the conversion of 5,10-methylene-tetrahydrofolate to 5-methyltetrahydrofolate (5THF), 5THF is the methyl donor for synthesis of methionine from homocysteinine [13]. The MTHFR gene is present on short arm of chromosome 1 at position p36.3. MTHFR A1298C polymorphism is associated with reduced MTHFR enzyme activity and hyperhomocysteinemia [5].

In A1298C polymorphism, A is substituted with C nucleotide at position 1298 [5], leading to substitution of glutamate by alanine s (Glu429Ala) in the MTHFR enzyme. Glu429Ala in MTHFR enzyme, reduces 40% enzyme activity. Frequency of 1298C allele differs greatly in various ethnic groups of the world. The prevalence of the mutant CC homozygote variant genotype ranges from 7 to 12% in Europe, 4 to 5% in Hispanics and 1 to 4% in Asian populations (1 to 4%) [14]. Several studies have reported A1298Cpolymorphism as risk factor for several diseases like-cleft lip and palate, Down syndrome, neural tube defects, and psychiatric disorders etc [14]. MTHFR polymorphisms were studied as risk for AD by several researcher but their results were controversial. Hence, the aim of present meta-analysis was to conclude the role of MTHFR A1298C polymorphism in AD risk.

## Methods

Article search was carried out in electronic databases up to August, 2019 using key terms - MTHFR’, ‘A1298C’, and ‘Alzheimer’s disease’. Criteria for inclusion of studies were as follows; (i) studies should be case-control association study; and (ii) the articles must report the sample size, distribution of alleles or genotypes for estimating the odds ratio (ORs) with 95% confidence interval (CIs). Studies were excluded if one of the following existed: (i) case-only studies, and (ii) editorial, case reports or reviews. Meta-analysis was carried out according to the method of Rai et al [15] (.2014). Publication bias was calculated according to the method of Egger et al. [16]. All statistical analysis was done by MetaAnayst.

## Results

Total ten studies [17-26] were found suitable for the inclusion in the meta-analysis. In incuded ten studies number of cases was 1067and number of contro was 1527. The lowest sample size was 43 [23] and highest sample size was 162 [20, 21] in included studies (Table 1). Total cases genotype percentage of AA, AC and CC was 55.34%, 44.65% and 11.84% respectively. The controls genotypes percentage of AA, AC and CC were 55.34%, 44.65% and 11.84% respectively.

**Table 1.**
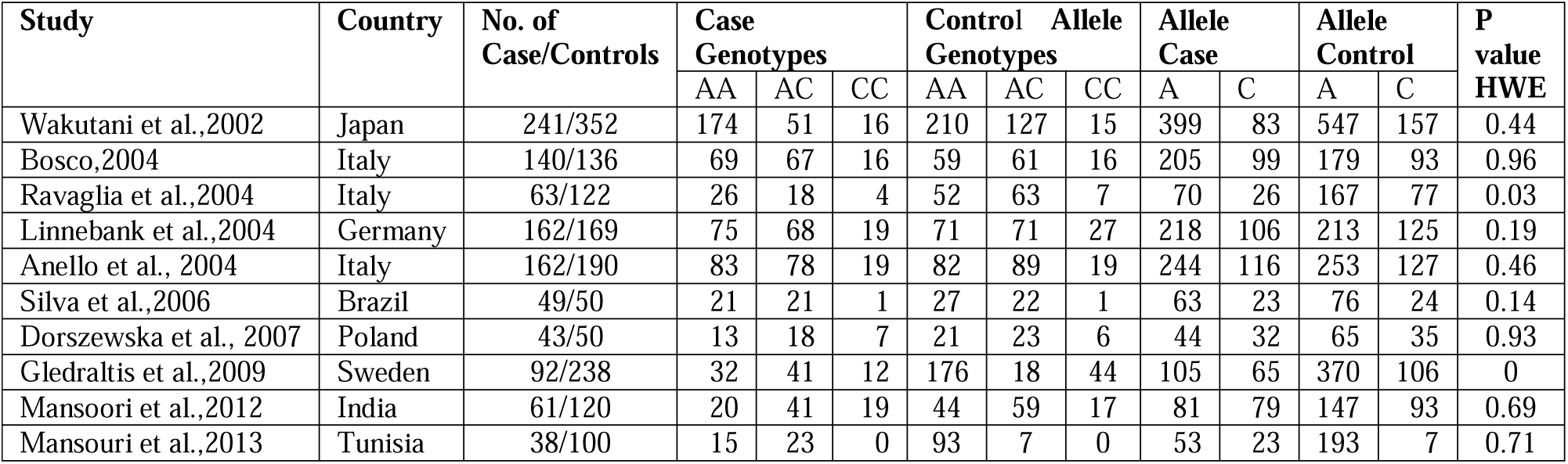
The distributions of MTHFR A1298C genotypes and allele number in Alzheimer’s disease cases and controls

In allele contrast (A vs C) meta-analysis, mutant C allele showed significant association with AD in random effect (OR= 1.26, 95%CI= 0.912-1.76, p= 0.04) models (Figure 1). Unlike to allele contrast meta-analysis, pooled odds ratio for homozygote genotype (CC vs. AA) did not show any association with AD adopting random (OR= 1.16, 95%CI= .88-1.55, p= 0.51) effect models. Association of mutant heterozygous genotype (AC vs. AA; co-dominant model) aso did not show any association with AD using random effect models (OR= 1.55; 95%CI= 0.81-2.93). Dominant mutant genotypes (CC+AC vs. AA) showed no association with AD using random (OR= 1.43; 95%CI= 0.85-2.44; p=0.05) effect models. Allele contrast cumulative meta-analysis showed that after inclusion of Gledraltis et al. [24] study, odds ratio increased to 1.029 and after then it increased to 1.26 (Figure 2). As evident by funnel plot the publication bias was not observed in any genetic model.

**Figure 1.**
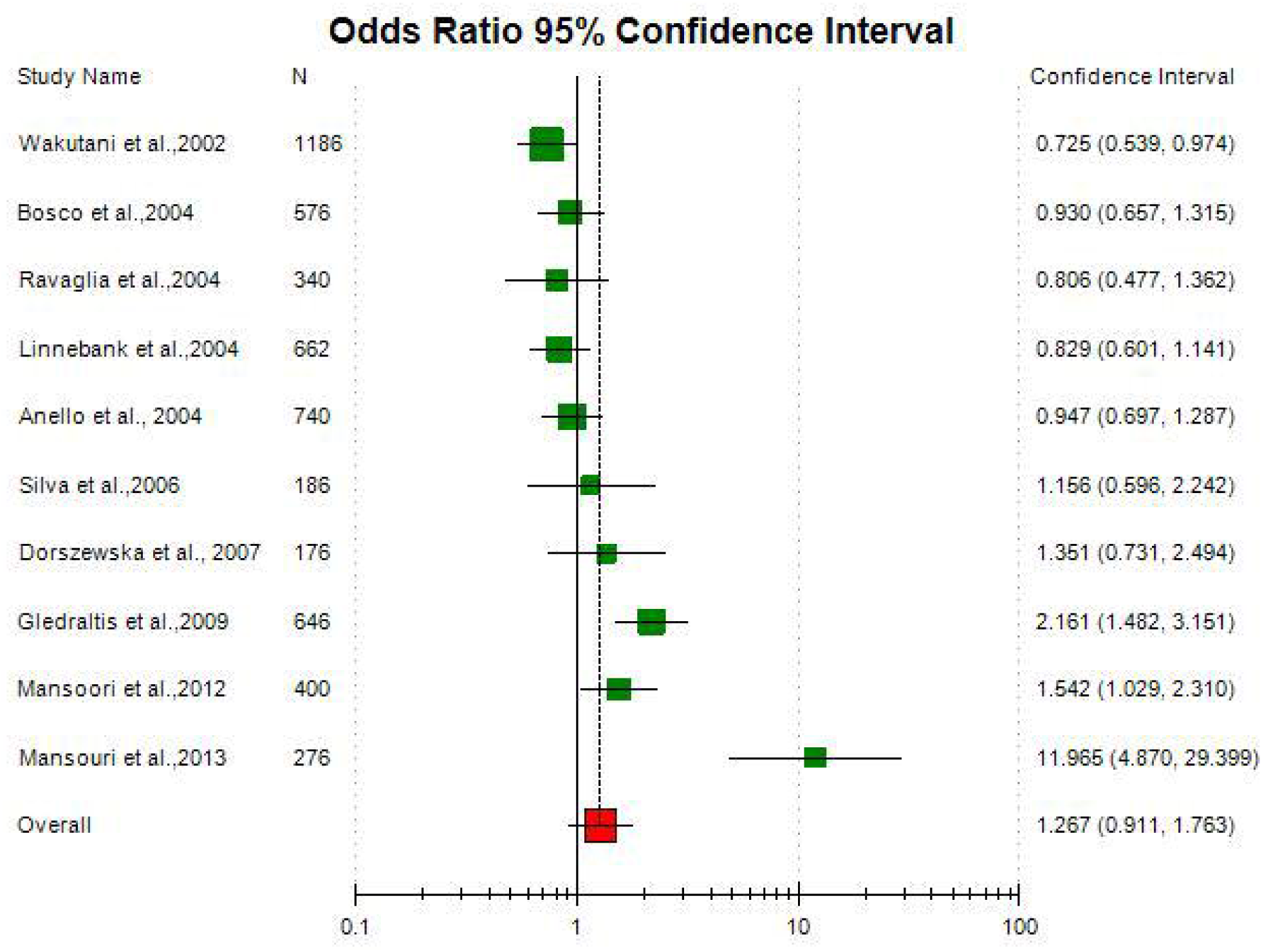
Allele Contrast (A vs C) Random Effect Forest Pot of Ten Studies.

**Figure 2.**
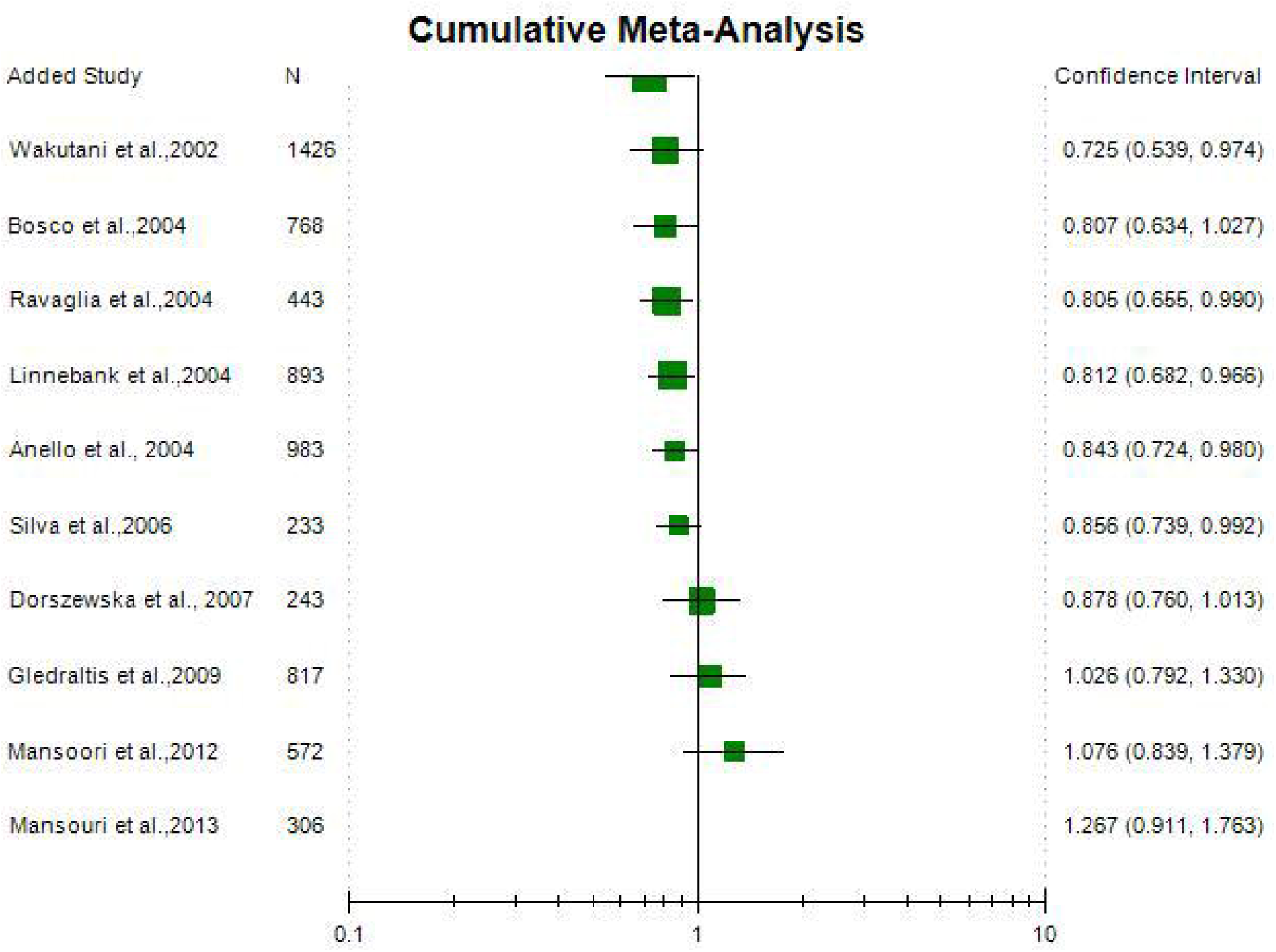
Allele Contrast Cumulative Meta-analysis.

## Discussion

Results of present meta-analysis showed no association between MTHFR A1298C polymorphism and Alzheimer’s disease. Elevated Hcy has been reported to be a risk factor for several psychiatric and neurodevelopmental disorders like neural tube defects, schizophrenia, bipolar disorder and depression. Hcy is implicated in increased oxidative stress, DNA damage, the triggering of apoptosis and excitotoxicity, all important mechanisms in neurodegeneration [27,28].

Meta-analysis is a statistical tool, which is successfully used for the compilation of contradictory results of small effect/power case-control studies. Several meta-analysis were published which evaluated risk of small effect gene polymorphism for different disease and disorders like-epilepsy [29], Alzheimer’s disease [30], G6PD [31], Down syndrome [32-34], Uterine Leiomyoma [35], orofacial cleft [36,37], depression [38], schizophrenia [39,40], autism [41,, male infertility [42] digestive tract cancer [43], lung cancer [44], endometrial cancer [45], breast cancer [46,47], prostate cancer [48], colorectal cancer [49], and esophageal cancer [50] etc.

Limitations of the study should be acknowledged like—(i) unadjusted crude OR is used, (ii) study number (only ten studies) and sample size are small, (iii) gene–gene or gene-environment interactions may modify the AD risk and; however, such stratified analysis could not be performed owing to lack of data.

